# Direct Whole-Blood PCR for Molecular Diagnostics

**DOI:** 10.64898/2026.11.10.765582

**Authors:** Jongwon Lim, Elyse Stachler, Stella Nielsen, Resham Talwar, Seungchan Baek, Kyle McMahon, Tsion Abay, Bryan Hsu, Dora Nabatanzi, Nallely Espinoza, Ruthie Mitchell, Hannah Knoll, Colby Wilkason, Al Ozonoff, Pardis Sabeti

## Abstract

Outbreaks of Ebola and other viral diseases underscore the need for rapid molecular diagnostics that can be performed directly from blood, yet conventional PCR workflows require sample processing that limits testing in remote settings. Here, we establish an extraction-free workflow for quantitative PCR (qPCR) and reverse-transcription qPCR directly in whole blood. Using 2 µL of blood per reaction, we combine blood-compatible amplification chemistry, RNase inhibition for RNA targets, and longer-wavelength fluorescence detection to address both biochemical and optical interference. We demonstrate detection of hepatitis B virus DNA at 10 copies/µL and RNA from hepatitis C, Bundibugyo, Ebola, and Sudan viruses at 100 copies/µL in blood spiked with viral nucleic acid standards. A portable instrument detected the Bundibugyo target with less than two minutes of hands-on preparation and results within one hour. These findings establish analytical feasibility for simplified blood-based molecular testing and motivate validation using intact viruses and clinical specimens.

## INTRODUCTION

Polymerase chain reaction (PCR) and reverse transcription PCR (RT-PCR) enable sensitive, specific detection and quantification of pathogen nucleic acids for infectious disease diagnosis.^1,2^ However, conventional blood-based PCR-based workflows typically rely on extensive pre-analytical processing, including sample fractionation, centrifugation, and multi-step nucleic acid extraction and purification, before amplification (**Fig. 1a**).^3^ Although these procedures isolate nucleic acids from complex biological matrices and improve downstream assay reliability, they increase turnaround time, hands-on complexity, and dependence on centralized laboratory infrastructure and trained personnel.^4^

**Figure 1.**
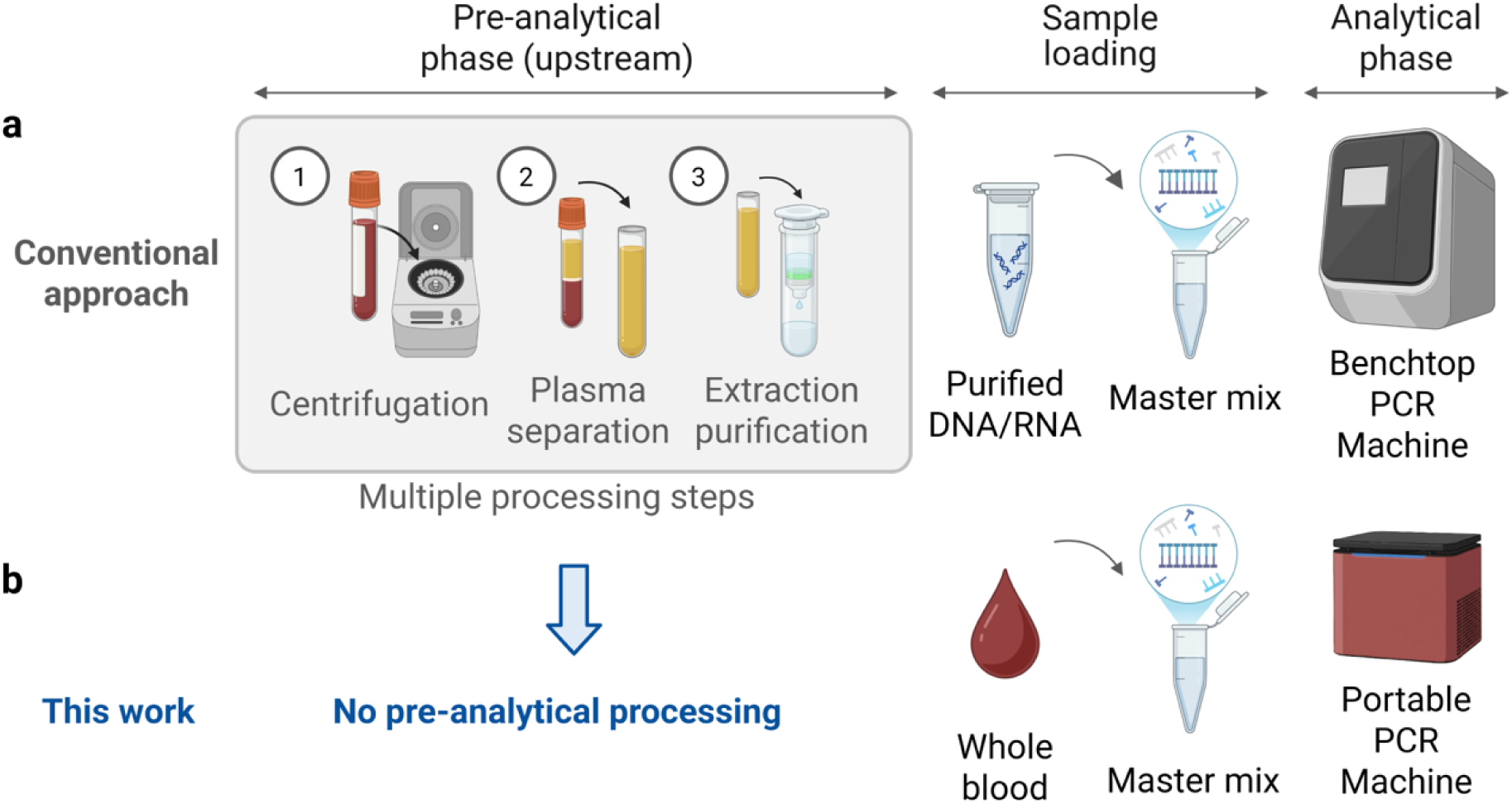
Comparison of conventional and direct whole-blood molecular diagnostic workflows. (A) Conventional molecular testing requires multiple pre-analytical processing steps, including centrifugation, plasma separation, and nucleic acid extraction and purification, before purified DNA or RNA is added to the reaction master mix for downstream analysis. (B) In this work, pre-analytical processing is eliminated by directly adding whole blood to the reaction master mix, enabling molecular analysis without centrifugation, plasma separation, or nucleic acid extraction and purification.

Constraints in molecular testing are particularly consequential when diagnostic speed or testing frequency directly affect clinical and public health decision-making at the point-of-care (POC).^5,6^ During acute infectious-disease outbreaks, including the COVID-19 pandemic,^7^ Mpox outbreak,^8^ and recent outbreaks caused by Bundibugyo virus (BDBV),^9,10^ rapid molecular detection is essential for timely triage, patient isolation, clinical management, and interruption of transmission. Similar logistical constraints affect chronic infections caused by hepatitis B virus (HBV) and hepatitis C virus (HCV), for which repeated measurements of viral load are important for disease assessment and evaluation of treatment response.^11,12^ Thus, reducing the pre-analytical burden associated with PCR-based testing remains an important step toward faster and more decentralized molecular diagnostics.^13,14^

POC nucleic acid amplification tests (NAATs) offer simplified molecular testing using PCR or isothermal amplification. These platforms integrate sample processing, amplification, and detection. Microfluidic cartridges,^15,16^ automated sample-processing modules,^17,18^ and self-contained diagnostic platforms^16^ can reduce manual intervention and shorten the distance between specimen collection and actionable molecular results. Isothermal amplification methods, including loop-mediated isothermal amplification (LAMP)^19,20^ and recombinase polymerase amplification (RPA),^21^ eliminate thermal cycling and, in many implementations, exhibit greater tolerance to crude biological matrices than conventional Taq polymerase-based PCR.^22^ Several isothermal platforms have advanced to POC diagnostics cleared or authorized by the U.S. Food and Drug Administration (FDA).^23,24^

Most successful POC NAATs, however, have focused on respiratory specimens, whereas direct whole-blood testing remains substantially less developed. These tests typically use relatively accessible specimens such as nasal or nasopharyngeal swabs and saliva. Whole blood is an information-rich and clinically important specimen for systemic infections, but it is also among the most challenging specimen matrices for NAATs.^25^ Target viral or microbial nucleic acids may be present at low abundance within a matrix dominated by erythrocytes, leukocytes, proteins, nucleases, heme-containing compounds, and other endogenous components that can interfere with amplification and optical detection.^3,26,27^

Direct PCR can bypass nucleic acid extraction, but existing approaches remain limited by sample dilution, additional processing, or reduced sensitivity.^28^ Most approaches rely on substantial sample dilution to mitigate blood-derived inhibition,^27,29–31^ improving amplification compatibility, but also reducing target concentration. While this approach may still be useful for high-abundance targets, it is particularly detrimental to low-abundance pathogens. In addition, direct PCR lacks the target-enrichment advantage of conventional extraction workflows due to its low sample input volume of a few microliters.^22^ Alternative strategies, including thermal^29,32^ or freeze–thaw pretreatment^33^ and nested PCR,^34^ add processing steps, prolong turnaround time, and may increase carryover contamination risk through repeated amplification or open-tube transfers. Inhibitor-tolerant or engineered polymerases have also been developed to enable amplification from blood, but analytical sensitivity may remain limited.^34,35^

Here, we set out to develop an extraction-free workflow for direct whole-blood qPCR and RT-qPCR that minimizes pre-analytical processing (Fig. 1b). We evaluate inhibitor-tolerant amplification chemistry and RNase inhibition to address the challenges of amplifying DNA and RNA directly from whole blood. We also investigate fluorescence detection at longer wavelengths as a strategy to reduce blood-associated optical interference. To assess the broader applicability of the approach, we evaluate analytical performance using whole blood spiked with HBV DNA and RNA from HCV, BDBV, Ebola virus (EBOV), and Sudan virus (SUDV). Finally, we examined whether the workflow can be transferred to a portable thermal cycler (Mic qPCR), with the goal of supporting decentralized testing for outbreak response and routine viral load monitoring.

## RESULTS

### Direct detection of HBV synthetic DNA from whole blood

To establish a benchmark for direct whole-blood testing, we first evaluated HBV synthetic DNA using standard qPCR in water detection. The baseline assay achieved a limit of detection (LOD) of 1 copy/µL with an amplification efficiency of 97.9%, calculated from the slope of the standard curve as E = (10^(−1/slope) − 1) × 100% (Fig. 2a and 2b).^36^

**Figure 2.**
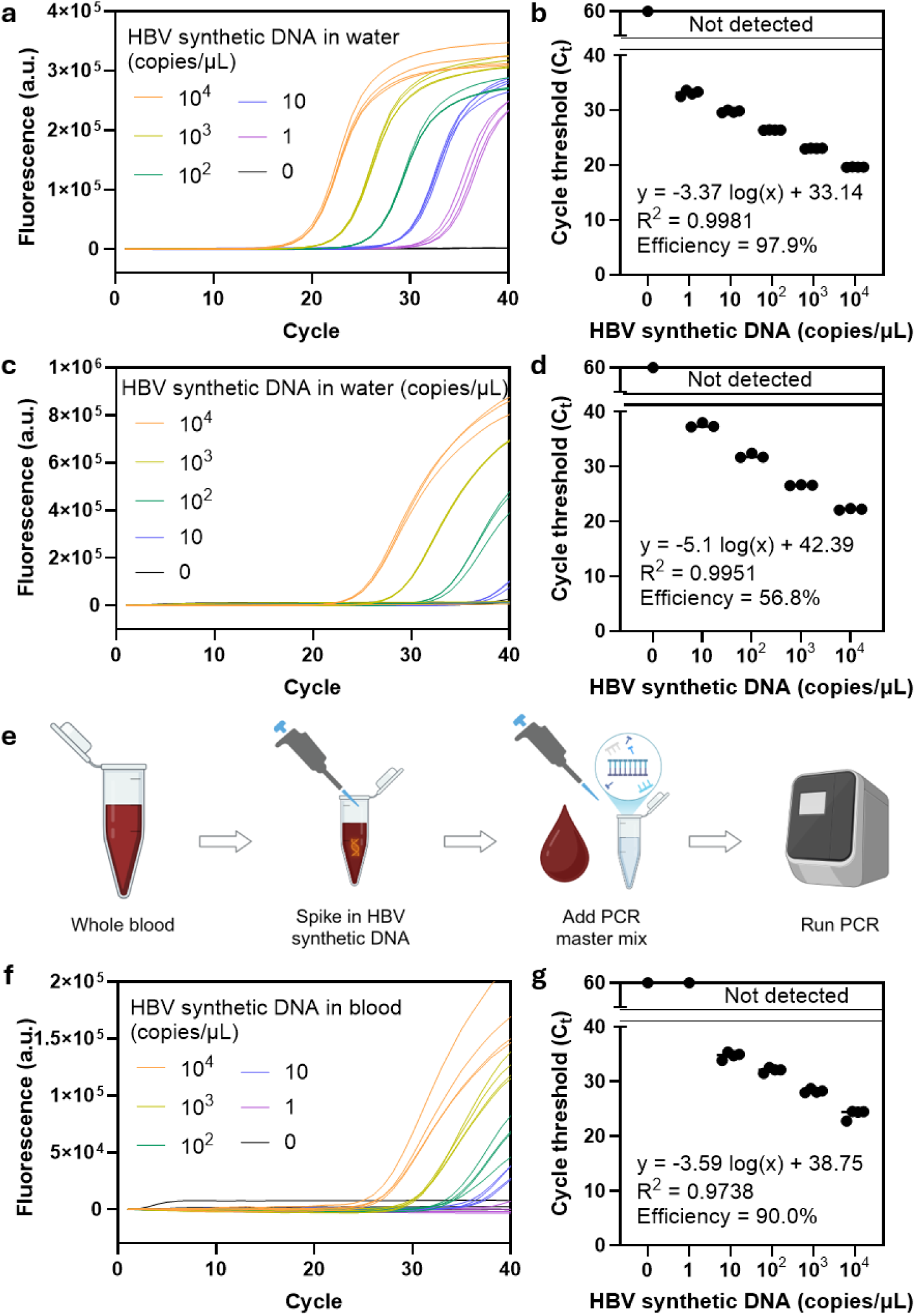
Direct detection of HBV synthetic DNA from whole blood using a blood-compatible PCR system. (A–D) Validation of HBV synthetic DNA amplification in water using conventional and blood-compatible PCR chemistries. Real-time qPCR amplification curves over 40 cycles are shown for PowerTrack™ SYBR Green Master Mix (A) and Q5® Blood Direct 2X Master Mix (C), with the corresponding cycle threshold (Ct) values summarized in (B) and (D), respectively. (E) Schematic of the extraction-free direct PCR workflow for HBV DNA detection. HBV synthetic DNA was spiked directly into whole blood, followed by addition of Q5 Blood Direct Master Mix and PCR amplification without any pre-analytical steps. (F–G) Direct detection of HBV synthetic DNA from whole blood using Q5 Blood Direct 2X Master Mix. Real-time qPCR amplification curves over 40 cycles are shown in (F), with the corresponding Ct values summarized in (G). Data are presented as individual points with mean (n = 3 to 4), and bars represent the mean.

We next tested a blood-compatible PCR formulation in water to determine how its performance differed from the standard assay before introducing blood. We selected Q5 Blood Direct Master Mix, a reagent formulated to tolerate up to 30% whole blood and anticoagulants such as sodium EDTA. Compared with standard qPCR in water, the Q5 assay had a higher LOD (10 versus 1 copies/µL) and lower estimated amplification efficiency (56.8% versus 97.9%); Fig. 2c and 2d). The lower performance in water may reflect the Q5 buffer formulation, which is optimized for complex crude matrices rather than purified aqueous targets.

We then tested whether the blood-compatible Q5 reagent could detect and quantify HBV DNA directly in whole blood without extraction. (Fig. 2e). We spiked HBV synthetic DNA directly into whole blood, performed serial dilutions, and amplified the samples directly in the Q5 master mix. The LOD remained at 10 copies/µL, the same as for Q5 in water, while estimated amplification efficiency increased from 56.8% to 90.0% (Fig. 2f and 2g). The standard curve maintained strong linearity across the tested concentration range (R² = 0.9738), supporting the potential for quantitative measurement of HBV DNA directly from whole blood. Although the direct whole-blood assay had a tenfold higher LOD than standard qPCR in water (10 vs. 1 copy/µL), it eliminated major pre-analytical processing bottlenecks by bypassing nucleic acid extraction and purification. Despite successful amplification, overall fluorescence yield in whole blood was approximately five-fold lower than in the Q5 reactions in water (Fig. 2c vs. 2f), consistent with blood-associated optical attenuation.

### Direct detection of HCV synthetic RNA from whole blood

Direct amplification of RNA from whole blood presents two additional challenges: degradation by endogenous RNases and the need for a compatible reverse transcriptase. To establish a baseline without blood-derived inhibition, we amplified HCV synthetic RNA in nuclease-free water using a standard master mix (PowerTrack) supplemented with WarmStart® RTx Reverse Transcriptase to enable one-step RT-qPCR without a separate cDNA synthesis step.^40^ The baseline assay achieved an LOD of 1 copy/µL with an estimated amplification efficiency of 105.9% based on the standard curve (Fig. 3a and 3b). Although late nonspecific amplification occurred in the no-target control, the substantial cycle threshold (Ct) separation between the 1-copy/µL and 0-copy/µL samples allowed clear analytical differentiation. Replacing the standard master mix with the blood-compatible Q5 Blood Direct Master Mix under the same aqueous conditions yielded an LOD of 10 copies/µL and an efficiency of 82.7% (Fig. 3c and 3d).

**Figure 3.**
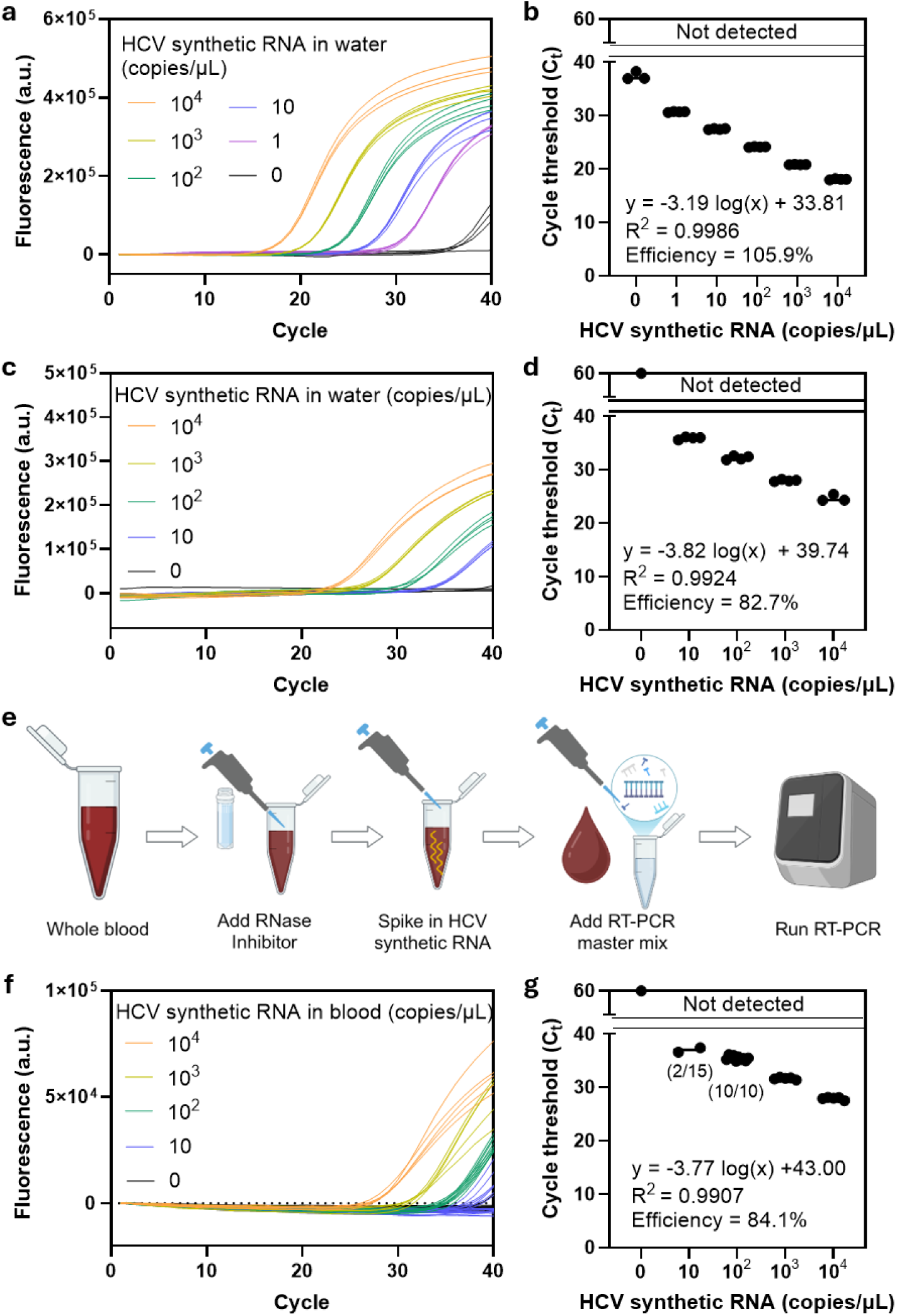
Direct detection of HCV synthetic RNA from whole blood using a blood-compatible RT-PCR system. (A–D) Validation of HCV synthetic RNA amplification in water using conventional and blood-compatible PCR chemistries supplemented with reverse transcriptase for RNA detection. Real-time RT-qPCR amplification curves over 40 cycles are shown for PowerTrack™ SYBR Green Master Mix supplemented with WarmStart RTx (A) and Q5® Blood Direct 2X Master Mix supplemented with WarmStart RTx (C), with the corresponding cycle threshold (Ct) values summarized in (B) and (D), respectively. (E) Schematic of the extraction-free direct RT-PCR workflow for HCV RNA detection. RNase inhibitor was first added to whole blood to minimize degradation of the RNA target by endogenous RNase activity, followed by spiking of HCV synthetic RNA and addition of Q5 Blood Direct Master Mix supplemented with reverse transcriptase, enabling direct RT-PCR without nucleic acid extraction or purification. (F–G) Direct detection of HCV synthetic RNA from whole blood using Q5 Blood Direct 2X Master Mix supplemented with Luna WarmStart RT Enzyme mix. Real-time RT-qPCR amplification curves over 40 cycles are shown in (F), with the corresponding Ct values summarized in (G). Data are presented as individual points with mean (n = 3 to 15), and bars represent the mean.

We added an RNase inhibitor to protect the HCV RNA target during direct whole-blood testing. We introduced the inhibitor directly into whole blood at a 1:39 volumetric ratio before spiking the sample with naked HCV synthetic RNA (Fig. 3e). Suppressing endogenous RNase activity is particularly important for analytical validation using naked viral RNA.^20^ In clinical specimens, by contrast, viral RNA is typically protected within intact viral particles prior to disruption.

We next evaluated reverse transcriptase compatibility and extension time to optimize amplification in the whole-blood matrix. We tested three enzymes: WarmStart® RTx Reverse Transcriptase, Luna® WarmStart® RT Enzyme Mix, and SuperScript™ IV Reverse Transcriptase. Using a reverse transcription (RT) incubation at 55°C for 10 minutes, we compared 15-second and 30-second extension protocols across HCV RNA concentrations of 10^3^, 10^2^, 10 and 0 copies/µL (Fig. S1). The 15-second extension paired with the Luna® WarmStart® RT Enzyme Mix delivered the best performance, demonstrating clear separation between target concentrations and high replicate-to-replicate consistency (Fig. S1b). In contrast, SuperScript IV produced substantial nonspecific amplification in the blood matrix (Fig. S1c).

We then assessed the analytical sensitivity and quantitative performance of the selected whole-blood RT-qPCR protocol. Using Q5 Blood Direct supplemented with Luna® WarmStart® RT and a 15-second extension, we detected 100 copies/µL in all replicates (10/10) and 10 copies/µL in a subset of replicates (2/15; Fig. 3f and 3g). The assay maintained a strong linear relationship between input RNA concentration and Ct (R² = 0.9907), with an estimated amplification efficiency of 84.1%. These findings support the feasibility of quantitative HCV RNA measurement directly from whole blood. As with the HBV DNA assay, maximum fluorescence yield was four- to five-fold lower in whole-blood than in Q5 reactions in water, consistent with blood-associated optical attenuation (Fig. 3c vs Fig. 3f).

The difference in analytical sensitivity between the DNA and RNA assays emerged primarily in whole blood. Under aqueous conditions, both qPCR and RT-qPCR achieved an LOD of 10 copies/µL when performed with Q5 Blood Direct Master Mix (Fig. 2d vs. Fig. 3d). In whole blood, however, direct qPCR maintained an LOD of 10 copies/µL for HBV DNA, whereas direct RT-qPCR reliably detected HCV RNA at 100 copies/µL, with only partial detection at 10 copies/µL (Fig. 2g vs. Fig. 3g). The minimal dilution introduced by the RNase inhibitor at a 1:39 volume ratio is unlikely to explain this difference. Unlike DNA qPCR, RT-qPCR requires an additional RT step, and the enzyme used here was not specifically optimized for activity in whole blood. Residual blood-associated inhibition of the RT reaction may therefore account, at least in part, for the reduced RNA detection sensitivity. These findings identify direct RT in whole blood as an important target for further optimization to improve the analytical sensitivity of direct whole-blood RT-qPCR.

### Extension of direct whole-blood RT-qPCR to multiple orthoebolaviruses

We next tested whether the optimized direct whole-blood RT-qPCR workflow could extend beyond HCV to other RNA viruses. We applied the assay framework established in Fig. 3e to three members of the *Orthoebolavirus* genus: BDBV, EBOV, and SUDV. We maintained the same whole-blood sample preparation and RT-qPCR workflow across all three targets, adjusting amplification conditions to accommodate the target-specific primer sets.^37^ Orthoebolavirus infections can produce high concentrations of circulating viral RNA during acute disease, making rapid detection directly from whole blood particularly relevant for outbreak diagnostics.^38,39^ We therefore evaluated each assay across a broad RNA concentration range, from 10^5^ to 10 copies/µL in whole blood.

The workflow supported direct amplification of all three viral RNA targets, although amplification efficiency varied among the assays. The BDBV assay exhibited an estimated amplification efficiency of 99.0% (Fig. 4a and 4b), compared with 88.8% for EBOV (Fig. 4c and 4d) and 79.0% for SUDV (Fig. 4e and 4f). Despite these differences, all three assays exhibited strong standard-curve linearity (R² = 0.9826, 0.9969, and 0.9816 for BDBV, EBOV, and SUDV, respectively), supporting the potential for quantitative measurement across multiple orthoebolavirus targets. All three assays achieved a LOD of 100 copies/µL. Together, these results show that the direct whole-blood RT-qPCR workflow can be adapted to multiple orthoebolavirus targets through target-specific primer and thermal-cycling optimization.

**Figure 4.**
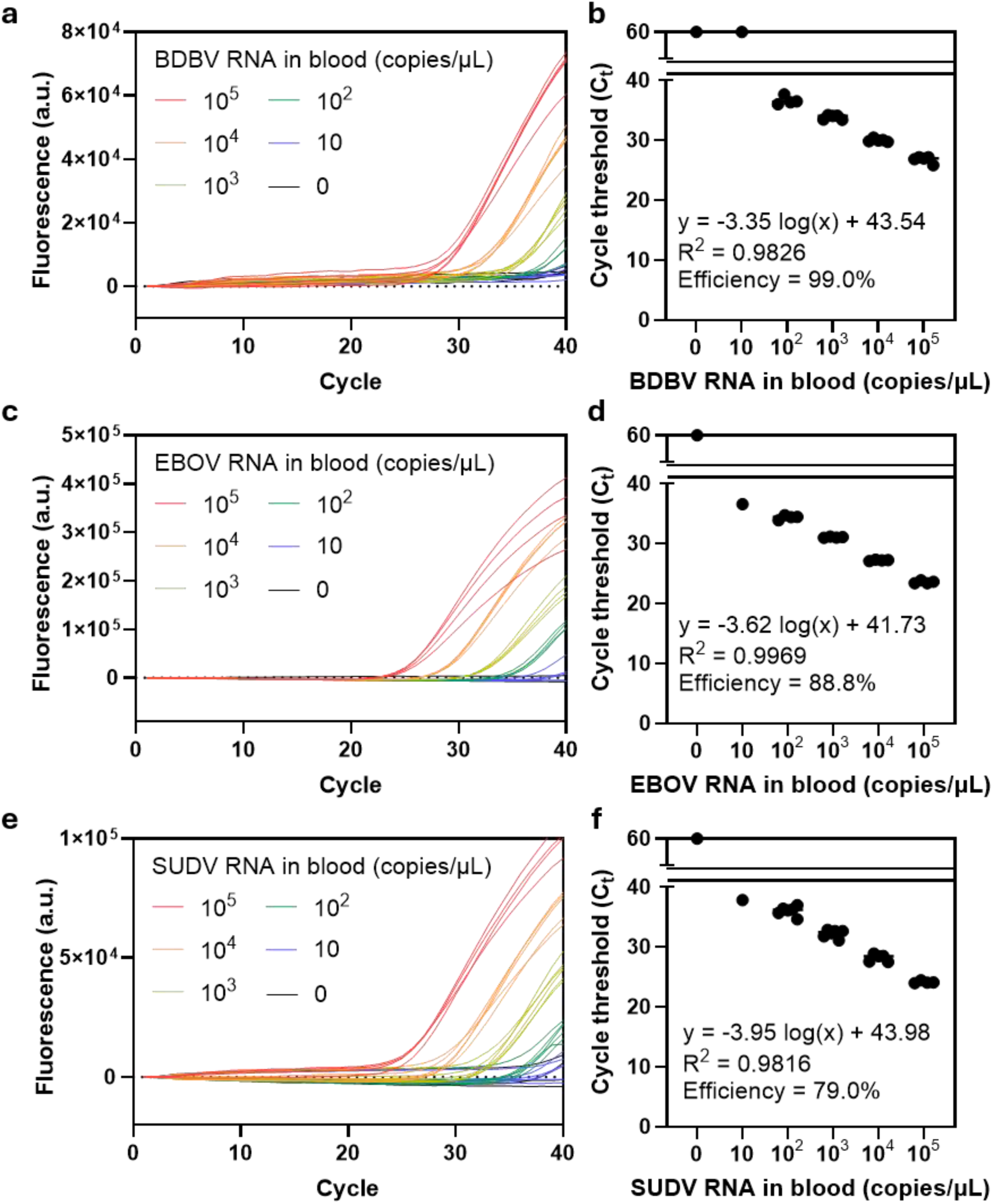
Extension of direct whole-blood molecular detection to multiple outbreak-relevant orthoebolaviruses. Direct whole-blood molecular detection was extended to three clinically important orthoebolaviruses: (A-B) BDBV, (C-D) Zaire ebolavirus (EBOV), and (E-F) Sudan virus (SUDV). Virus-specific assays were developed and evaluated to establish robust amplification for each target and subsequently assess their compatibility with the direct whole-blood detection workflow. Amplification curves demonstrate successful detection across all three viral targets, supporting the broader applicability of the direct detection platform beyond HBV and HCV and its potential adaptation to emerging and outbreak-associated viral pathogens. Data are presented as individual points with mean (n = 4 to 6), and bars represent the mean.

### Wavelength-dependent fluorescence attenuation

Whole blood can weaken the fluorescence signal used to detect amplification, even when the reaction is successful.^40,41^ This attenuation arises in part from heme- and hemoglobin-associated absorption and quenching,^42^ which are particularly pronounced at shorter visible wavelengths and decrease toward the red region of the spectrum (Fig. 5a). We therefore hypothesized that shifting detection toward longer excitation and emission wavelengths could improve fluorescence signal retention in the whole-blood matrix.

**Figure 5.**
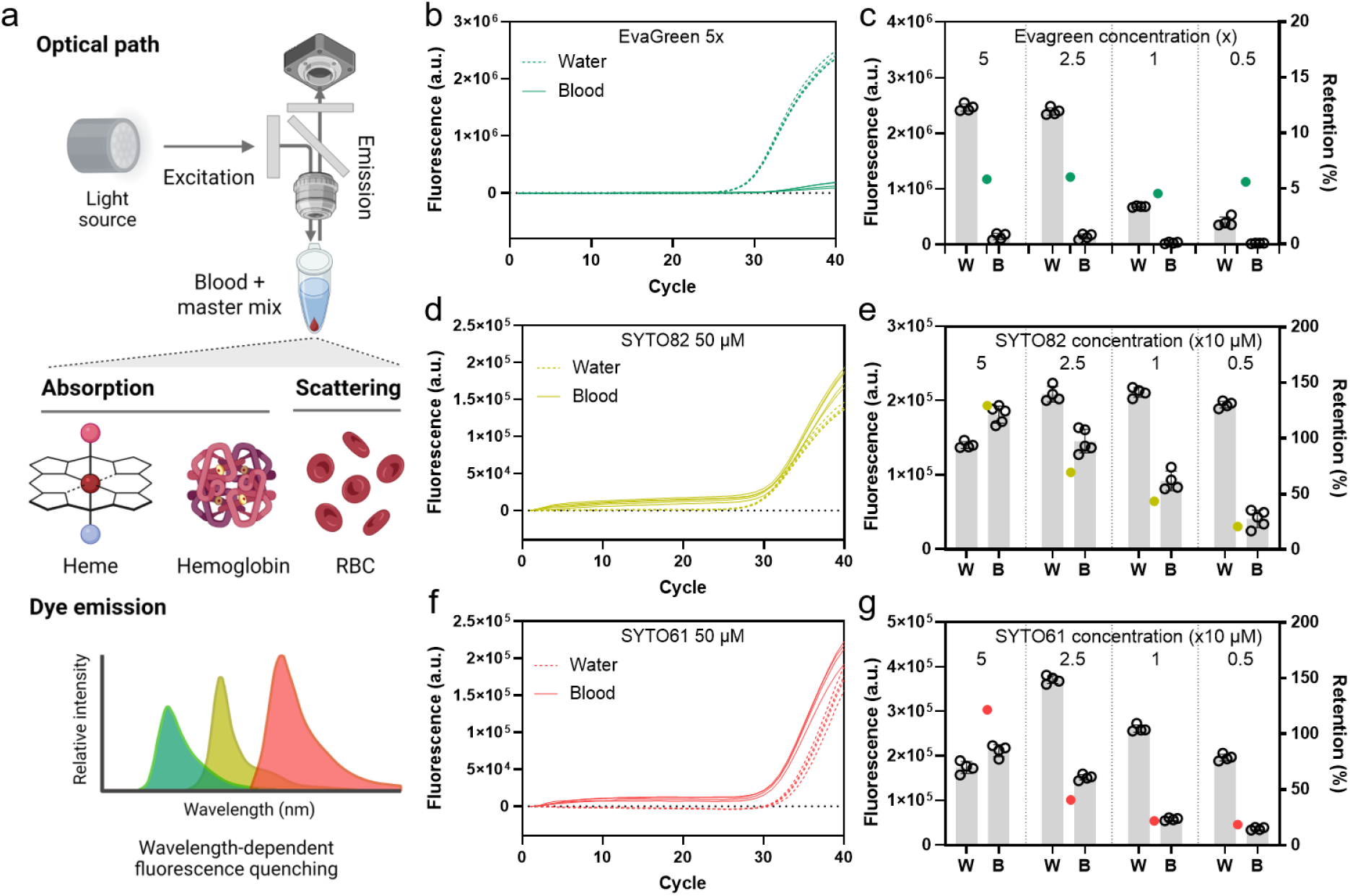
Wavelength-dependent fluorescence attenuation by whole blood during direct PCR with HBV detection protocol. (A) Schematic illustrating wavelength-dependent absorption and quenching by heme and hemoglobin, and light scattering by red blood cells. (B–G) Representative amplification curves and corresponding fluorescence intensity and retention measurements for EvaGreen (B, C; 0.5×–5×), SYTO 82 (D, E; 5–50 µM), and SYTO 61 (F, G; 5–50 µM) in water and whole blood using the Q5 Blood Direct HBV assay. Fluorescence retention (%) was calculated relative to the corresponding water control and is indicated by red dots in (C, E, G). Individual data points represent fluorescence measurements (n = 4–5), and gray bars indicate mean fluorescence intensity.

To test whether reporter wavelength affected fluorescence retention, we compared three intercalating dyes using the Q5 Blood Direct HBV assay established in Fig. 2e. The dyes spanned three detection channels: EvaGreen (green), SYTO 82 (yellow/orange), and SYTO 61 (red). For each dye concentration, we expressed fluorescence intensity in successfully amplified whole-blood reactions as a percentage of that in the corresponding water control (fluorescence retention: Fig. 5b–g and Fig. S2). EvaGreen exhibited severe signal attenuation across concentrations ranging from 0.5× to 5× the manufacturer-recommended concentration. Whole-blood reactions retained less than 10% of the corresponding aqueous signal at every EvaGreen concentration tested, indicating poor signal retention with this green-channel reporter (Fig. 5b and 5c).

Fluorescence retention improved with the longer-wavelength reporter SYTO 82, although the degree of recovery depended on dye concentration. Across 5–50 µM, whole-blood reactions retained approximately 20–129% of the corresponding aqueous fluorescence signal (Fig. 5d and 5e). In water, SYTO 82 produced comparable fluorescence intensities at 5, 10, and 25 µM. In contrast, increasing the SYTO 82 concentration to 50 µM in whole blood produced fluorescence intensities comparable to, and in some measurements exceeding, the corresponding water control, with overlapping amplification curves and no noticeable delay relative to water.

The red-channel reporter SYTO 61 also improved fluorescence retention relative to EvaGreen. Across 5–50 µM, SYTO 61 retained approximately 18%-120% of the corresponding aqueous fluorescence signal (Fig. 5f and 5g). Fluorescence signals in water peaked at 25 µM, where whole-blood reactions retained approximately 40% of the aqueous signal. Although the degree of signal recovery varied with dye concentration, SYTO 61 consistently outperformed the green-channel EvaGreen readout. Together, these results support selecting longer-wavelength reporters and optimizing their concentrations as a practical strategy to improve fluorescence readout in direct whole-blood qPCR and RT-qPCR.

### Direct POC RT-qPCR

We next evaluated whether direct whole-blood RT-qPCR workflow could be transferred from a benchtop instrument to the portable Mic qPCR system (Fig. 6a). The simplified workflow consisted of four steps: (1) collection of a small volume of whole blood, compatible with finger-prick sampling; (2) addition of RNase inhibitor to the blood sample; (3) addition of the treated sample and RT-qPCR master mix to Mic-compatible reaction tubes; and (4) loading of the tubes into the portable instrument for amplification and fluorescence detection. The sample-to-instrument workflow required less than 2 minutes of hands-on time, followed by 55 minutes of automated RT-qPCR. The total sample-to-result was less than 1 hour, without nucleic acid extraction or purification.

**Figure 6.**
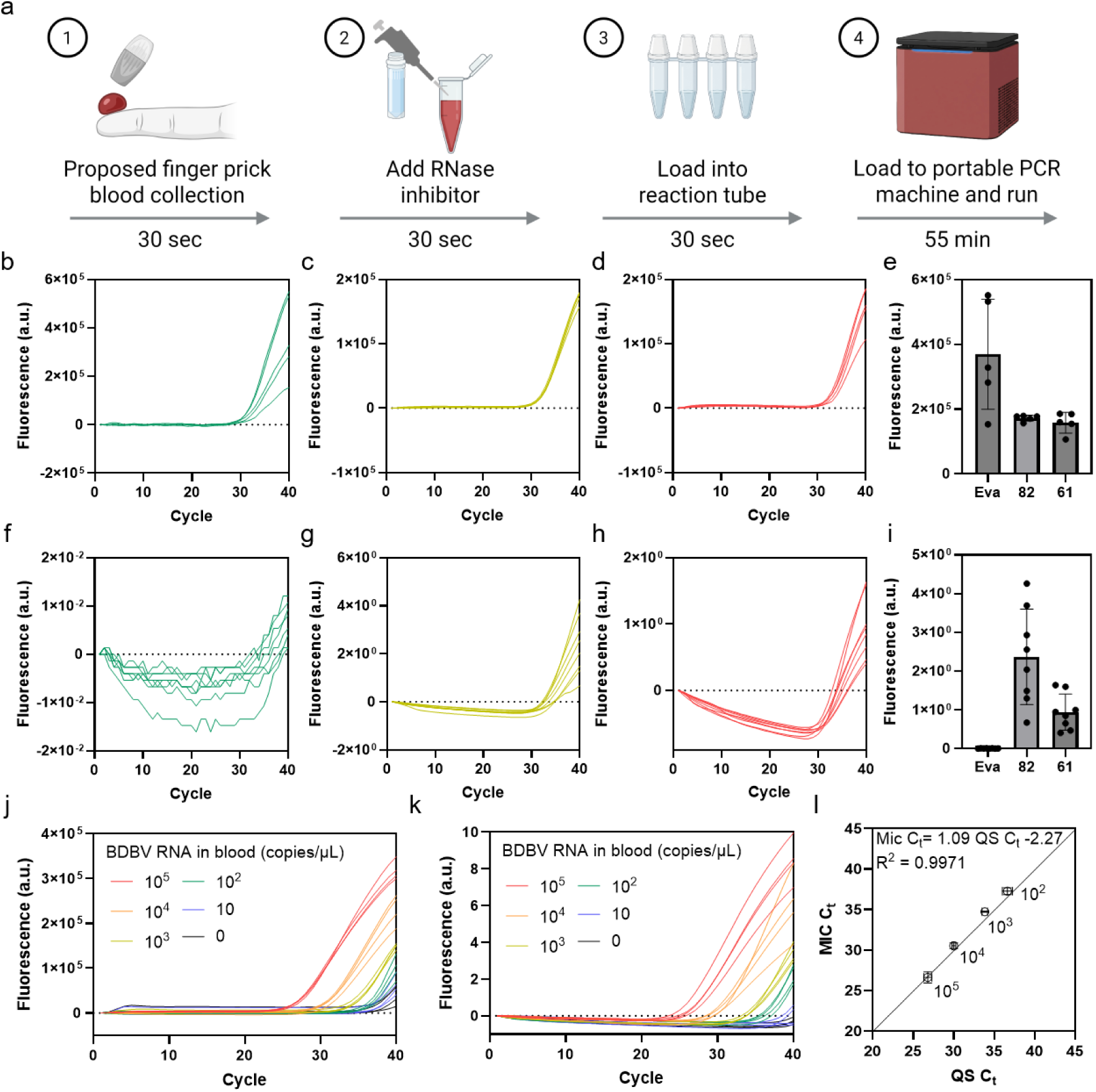
Translation of direct whole-blood RT-qPCR to a portable PCR platform. (A) Schematic of the four-step workflow comprising finger-prick blood collection (proposed, 30 s), RNase inhibitor addition (30 s), reaction preparation (30 s), and automated RT-qPCR on the portable Mic system (55 min), enabling sample-to-result detection in under 1 h without nucleic acid extraction. (B–I) Comparison of HBV DNA detection (10² copies/µL) in whole blood using the Q5 Blood Direct assay on the QuantStudio (B–E) and Mic (F–I) systems. Representative amplification curves are shown for EvaGreen (5×, green channel; B, F), SYTO 82 (50 µM, yellow channel; C, G), and SYTO 61 (50 µM, red channel; D, H), with corresponding endpoint fluorescence intensities summarized in (E, I). (J, K) Direct whole-blood RT-qPCR detection of BDBV RNA across serial concentrations using SYTO 82 (50 µM) on the QuantStudio (J) and Mic (K) systems. (L) Correlation of Ct values between the Mic and QuantStudio systems for BDBV detection, showing a linear regression slope of 1.09 and R² = 0.9971. Individual fluorescence measurements are shown for panels B–I (n = 5–8), with gray bars indicating mean endpoint fluorescence intensities in (E, I). Individual amplification curves are shown for (J, K) (n = 4), and data in (L) are presented as mean ± SD.

Because fluorescence detection characteristics differ among real-time PCR instruments,^43,44^ we tested whether the fluorescence advantage of longer-wavelength reporters persisted on the Mic system. Using the Q5 Blood Direct HBV assay, we compared three reporter conditions (Fig. 6b–i): EvaGreen at 5× concentration in the green channel (Fig. 6b and 6f), SYTO 82 at 50 µM in the yellow channel (Fig. 6c and 6g), and SYTO 61 at 50 µM in the red channel (Fig. 6d and 6h). On the benchtop QuantStudio instrument (Fig. 6b-d), all three reporter conditions enabled detection of 10² copies/µL of HBV DNA directly in whole blood, with endpoint fluorescence intensities within the same order of magnitude (approximately 1.5–6 × 10⁵ arbitrary fluorescence units; Fig. 6e).

Differences in reporter signal intensity were more pronounced on the Mic system than on QuantStudio (Fig. 6f-i). We observed strong amplification-associated fluorescence in the yellow and red channels, whereas the green-channel signal remained markedly weaker despite a small increase during amplification (Fig. 6f–i). The green-channel signal was two orders of magnitude lower than the red-channel signal and several hundred-fold lower than the yellow-channel signal under the tested conditions.

Reporter performance in whole blood therefore depends on both intrinsic spectral properties of the fluorophore and instrument optics, including channel-specific gains and sensitivity.^43–45^ Despite these instrument-specific differences, the longer-wavelength reporters provided stronger amplification-associated signals than the green-channel reporter on the Mic system, consistent with the wavelength-dependent effects observed in Fig. 5. Thus, empirical selection of fluorophore and detection channel is particularly important when transferring direct whole-blood assays between instruments. Among the conditions tested, SYTO 82 in the Mic yellow channel provided the strongest signal, and we selected it for subsequent portable RT-qPCR experiments.

We next evaluated portable RNA detection using SYTO 82 and direct whole-blood BDBV RT-qPCR assay established in Fig. 4a and 4b. Using SYTO 82 as the reporter, the BDBV assay detected 100 copies/µL on the QuantStudio platform, although low-level nonspecific amplification was observed in the 0-copy/µL control under these conditions (Fig. 6j). On the Mic system, BDBV RNA was detected in all replicates at 100 copies/µL (4/4; Ct 36.8–37.8). At 10 copies/µL, no reaction met the detection criterion (Ct ≤ 38.0): two of four reactions showed only a late rise that crossed the threshold at Ct ≈ 39.8, and the other two did not cross the threshold by cycle 40. None of the 0-copy/µL controls crossed the threshold (0/4; Fig. 6k). These results demonstrate that direct whole-blood BDBV RT-qPCR can be performed on the portable platform while retaining analytical sensitivity in the 100 copies/µL range.

Finally, we compared BDBV Ct measurements to assess the consistency of results between the benchtop and portable system. We compared values from the Mic/SYTO 82 configuration with those from the QuantStudio calibration curve for the BDBV assay (Fig. 4b). Ct measurements were strongly correlated between the two platforms, with a near-unity regression slope of 1.09 and R² = 0.9971 (Fig. 6l). Together, these results demonstrate that direct whole-blood RT-qPCR can be transferred to a portable real-time PCR platform by matching reporter wavelength to instrument-specific optical performance. The combination of minimal sample handling, extraction-free amplification, long-wavelength fluorescence detection, and portable instrumentation provides a practical framework for decentralized molecular detection of blood-borne viral targets.

## DISCUSSION

We developed a direct whole-blood qPCR and RT-qPCR workflow that detects DNA and RNA viral targets without centrifugation, nucleic acid extraction, or purification. Using 2 µL of whole blood spiked with viral nucleic acids, we detected HBV DNA, and RNA from HCV, BDBV, EBOV, and SUDV at 10–100 copies/µL, showing that the approach is not limited to a single target or nucleic acid class. On a portable platform, direct whole-blood BDBV RT-qPCR required less than 2 min of hands-on preparation and less than 1 h from sample to result. Its Ct values correlated closely with those from the benchtop system.

Our simplified workflow enabled direct detection of both DNA and RNA targets from whole blood, but analytical sensitivity could be further improved. The direct assays were less sensitive than standard assays tested in water, and HCV RNA detection was less sensitive than HBV DNA detection in whole blood. For HBV, the blood-compatible formulation detected 10 copies/µL in both water and whole blood, compared with 1 copy/µL for standard qPCR in water. This sensitivity difference was therefore already present before blood was introduced. For HCV, however, the blood-compatible assay detected 10 copies/µL in water, whereas detection in whole blood was consistent at 100 copies/µL but only partial at 10 copies/µL. Residual inhibition of reverse transcription may contribute to the reduced RNA sensitivity, although the experiments do not isolate its effect from other differences between assay conditions. Further work to distinguish RNA degradation from inefficient reverse transcription could help guide improvements.

Whole blood attenuates fluorescence, and longer-wavelength reporters mitigated this effect. This improvement offers a practical way to strengthen fluorescence readout without adding sample-processing steps (Fig. 5). The best reporter also depended on the instrument: on the Mic system, SYTO 82 in the yellow channel gave the strongest signal and the green channel the weakest (Fig. 6). These findings highlight the need to select the fluorescent dye and detection channel together when adapting an assay to a new instrument. Addressing amplification chemistry and optical detection together is therefore important for developing reliable direct whole-blood tests.

The workflow has practical strengths within the REASSURED framework, which places analytical performance alongside the practical requirements of POC testing, including affordability, ease of use, and speed (Table S1). These strengths include a small blood-volume requirement, minimal sample processing, and less than 2 minutes of hands-on preparation. Avoiding extraction reduces the processing steps and consumables needed between sample collection and amplification. The small sample input could also support finger-prick collection, although we did not evaluate finger-stick specimens. The estimated reagent and consumable cost of approximately $2.53 per reaction suggests potential for affordable testing, although this estimate excludes instrument and labor costs and does not represent the full cost of a clinical test (Table S2). Together, these features could make testing near the site of collection more practical and reduce dependence on specimen transport to centralized laboratories (Supplementary Note 1).

Direct blood assays differ widely in the sample processing they require, from pretreated or diluted specimens to minimally manipulated whole blood.^46^ Previous approaches have used additional analytical steps for direct dengue detection, repeated freeze–thaw pretreatment for Ebola detection, or two rounds of amplification for direct blood RT-PCR (Table S3).^27,47,48^ A direct whole-blood Zika RT-qPCR assay also avoided additional processing, providing an important precedent for this approach, however with approximately fivefold higher (9.5 x 10^2^ copies/reaction) LOD than ours.^49^ Cartridge-based systems such as GeneXpert Ebola report a lower detection limits (∼230 copies/mL) but depend on complex microfluidic consumables (Supplementary Note 2).^50^ These approaches offer different balances of sensitivity, sample input, cost, and processing complexity. Extraction can concentrate targets from a larger specimen, whereas direct testing removes those processing steps but analyzes a small volume of blood. Comparisons should therefore consider both the concentration in the original specimen and the amount of that specimen ultimately represented in the amplification reaction.

Several limitations remain in analytical performance, validation, and practical implementation. Inconsistent detection at low concentrations and nonspecific amplification under some conditions warrant further refinement of sensitivity, specificity, and result interpretation. Previous work reported no cross-reactivity of the BDBV assay with EBOV or SUDV and no false-positive amplification in healthy human plasma,^51^ but specificity requires separate validation for the direct whole-blood workflow evaluated here. Quantitative accuracy and reproducibility also require validation. Our experiments used synthetic or purified nucleic acids rather than intact viruses or patient specimens, leaving detection of nucleic acids within viral particles and clinical performance untested. Broader evaluation should include different blood specimens and collection conditions, including finger-prick samples, and additional RNA targets on the portable platform, where RNA testing was limited to BDBV. The workflow also uses liquid reagents and external power and requires separate analysis of exported fluorescence data. Reagent stabilization, integrated battery operation, and automated result interpretation could simplify use outside the laboratory (Supplementary Note 1). Evaluation in the intended setting will be important to establish complete turnaround time, ease of use, and performance under routine operating conditions.

Together, these findings provide a foundation for simpler blood-based molecular testing by combining minimal sample preparation with blood-compatible amplification and fluorescence detection. Further refinement and validation could support testing closer to the site of sample collection, with potential applications in outbreak response and viral load monitoring.

## MATERIALS AND METHODS

### Whole-blood samples, nucleic acids and primers

Human whole blood was purchased from Innovative Research (Cat. No. IWB1NAE10MLC) and stored at 4 °C with continuous rotation to prevent coagulation. Quantitative synthetic hepatitis B virus (HBV) DNA (ATCC, VR-3232SD) and hepatitis C virus (HCV) RNA (ATCC, VR-3233SD) were aliquoted upon receipt and stored at −80 °C. Synthetic DNA templates containing the target regions of Bundibugyo virus (BDBV), Ebola virus (EBOV), and Sudan virus (SUDV) were obtained from Twist Bioscience in lyophilized form. Following resuspension, RNA templates were generated by in vitro transcription using the HiScribe T7 High Yield RNA Synthesis Kit (New England Biolabs [NEB], E2040L) according to the manufacturer’s instructions. Residual DNA was removed by treatment with RNase-free DNase I (NEB, M0303S), and the RNA was purified using RNAClean XP beads (Beckman Coulter, A63987). Purified RNA was quantified using the Qubit RNA High Sensitivity Assay Kit (Invitrogen, Q32852), normalized to 10⁸ copies/µL, aliquoted, and stored at −80 °C. Individual aliquots were used for each experiment and were not reused. Primer sequences were adapted from previously published assays and synthesized by IDT. All primer sequences and their corresponding references are provided in Table S4.

### Reagents

PCR amplification was performed using PowerTrack SYBR Green Master Mix (Applied Biosystems, A46109) or Q5 Blood Direct 2X Master Mix (NEB, M0500L). Reverse transcription was evaluated using WarmStart RTx Reverse Transcriptase (NEB, M0380L), Luna WarmStart RT Enzyme Mix supplied with the Luna Universal One-Step RT-qPCR Kit (NEB, E3005L), and SuperScript IV Reverse Transcriptase (Invitrogen, 18090050). Murine RNase Inhibitor (NEB, M0314L) was used to protect RNA templates during direct whole-blood RT-qPCR. Fluorescence detection was performed using EvaGreen Dye, 20X in Water (Biotium, 31000), SYTO 82 Orange Fluorescent Nucleic Acid Stain (Invitrogen, S11363), or SYTO 61 Red Fluorescent Nucleic Acid Stain (Invitrogen, S11343).

### Standard qPCR and RT-qPCR in nuclease free water

Standard qPCR reactions were performed in a total volume of 20 µL by combining 15 µL of reaction mixture with 5 µL of serially diluted DNA template in nuclease-free water. Each reaction contained 1× PowerTrack SYBR Green Master Mix and 500 nM each forward and reverse primer. For RT-qPCR, the reaction mixture was additionally supplemented with 0.3 U of WarmStart RTx Reverse Transcriptase per reaction, and serially diluted RNA was used as the template. Unless otherwise indicated, reagent concentrations refer to final concentrations in the complete 20-µL reaction. Amplification was performed on a QuantStudio 6 Flex Real-Time PCR System (Applied Biosystems). The qPCR program consisted of polymerase activation at 95 °C for 2 min, followed by 40 cycles of denaturation at 95 °C for 5 s and annealing/extension at 60 °C for 30 s. For RT-qPCR, reverse transcription was performed at 55 °C for 10 min before polymerase activation. Ct values were determined automatically using the QuantStudio software and used to construct standard curves.

### Direct whole-blood qPCR for DNA detection

Direct whole-blood qPCR reactions were performed in a total volume of 20 µL by combining 18 µL of reaction mixture with 2 µL of whole blood containing serially diluted DNA template. Each reaction contained 1× Q5 Blood Direct Master Mix, 500 nM each forward and reverse primer, and 1× EvaGreen, unless otherwise specified. The whole-blood input represented 10% of the total reaction volume. Amplification was performed on the QuantStudio 6 Flex system using an initial denaturation at 98 °C for 3 min, followed by 40 cycles of denaturation at 98 °C for 1 s, annealing at the primer-specific temperature for 5 s, and extension at 72 °C for 15 s. Annealing temperatures were selected using the NEB Tm Calculator. An annealing temperature of 67 °C was used for the HBV assay. Because the reduced fluorescence intensity in whole-blood reactions prevented reliable automatic Ct determination under the tested conditions, raw fluorescence data were exported and analyzed using the procedure described under “Analytical performance and data analysis”.

### Direct whole-blood RT-qPCR for RNA detection

Murine RNase Inhibitor was mixed with whole blood at an inhibitor-to-blood volumetric ratio of 1:39 before RNA addition. RNA templates were serially diluted in the inhibitor-treated whole blood to obtain the desired concentrations. A 2-µL aliquot of each sample was then combined with 18 µL of reaction mixture. The reaction composition was the same as that used for direct whole-blood qPCR, with the addition of a reverse transcriptase. The enzymes evaluated were Luna WarmStart RT Enzyme Mix at 1×, WarmStart RTx Reverse Transcriptase at 0.3 U/µL, and SuperScript IV Reverse Transcriptase at 8 U/µL. Unless otherwise specified, Luna WarmStart RT Enzyme Mix was used for subsequent direct whole-blood RT-qPCR experiments. The thermal program consisted of reverse transcription at 55 °C for 10 min, initial denaturation at 98 °C for 3 min, and 40 cycles of denaturation at 98 °C for 1 s, annealing at the target-specific temperature for 5 s, and extension at 72 °C for 15 s. Annealing temperatures were 65 °C for HCV, 66 °C for BDBV, 63 °C for EBOV, and 64 °C for SUDV. Raw fluorescence data were analyzed as described below.

### Optimization of reverse transcriptase and extension time

Reverse transcriptase compatibility and PCR extension time were evaluated using HCV RNA at 10³, 10², 10, and 0 copies/µL in inhibitor-treated whole blood. All reactions included an initial reverse-transcription incubation at 55 °C for 10 min. WarmStart RTx Reverse Transcriptase, Luna WarmStart RT Enzyme Mix, and SuperScript IV Reverse Transcriptase were compared using a 15-s extension at 72 °C. WarmStart RTx and Luna WarmStart RT Enzyme Mix were additionally evaluated using a 30-s extension. All other reaction components and cycling conditions were maintained as described above. Performance was assessed by examining concentration-dependent separation of amplification curves, consistency among replicate reactions, and amplification in no-target controls. Each condition was tested using four to five replicate reactions.

### Evaluation of wavelength-dependent fluorescence attenuation

The effect of fluorescent reporter selection on signal retention in whole blood was evaluated using the Q5 Blood Direct HBV qPCR assay. EvaGreen was tested at 0.5×–5× the manufacturer-recommended working concentration. SYTO 82 and SYTO 61 were each tested at final concentrations of 5, 10, 25, and 50 µM. For each reporter concentration, amplification was performed in whole blood and in a corresponding nuclease-free water control. EvaGreen, SYTO 82, and SYTO 61 fluorescence was acquired using the QuantStudio SYBR, VIC, and Cy5 channels, respectively. Fluorescence intensities from successfully amplified reactions were compared between the two matrices. Fluorescence retention was calculated as: Fluorescence retention (%) = (fluorescence intensity in whole blood / fluorescence intensity in the corresponding water control) × 100. Endpoint fluorescence intensities were used to compare signal retention between whole-blood reactions and the corresponding water controls. Each condition was tested using four to five replicate reactions.

### Portable direct whole-blood PCR/RT-qPCR using the Mic system

Direct whole-blood qPCR and RT-qPCR were transferred to the Mic real-time PCR system using Mic-compatible reaction tubes. Reaction mixture and whole-blood sample were dispensed onto the inner wall of each tube. Tubes were then capped and loaded into the instrument. No further manual intervention was required during amplification. EvaGreen, SYTO 82, and SYTO 61 were detected using the Mic green, yellow, and red channels, respectively. Reporter performance was compared using the direct whole-blood HBV qPCR assay with 5× EvaGreen, 50 µM SYTO 82, or 50 µM SYTO 61. SYTO 82 at 50 µM was selected for subsequent portable BDBV RT-qPCR experiments. Reactions were performed using the target-specific thermal programs described above. Following amplification, raw fluorescence data were exported from the Mic software and analyzed using the same procedure applied to whole-blood reactions on the QuantStudio system.

### Analytical performance and data analysis

For reactions performed in nuclease-free water, Ct values were determined using the QuantStudio software with automatic baseline and threshold settings. For whole-blood reactions, low fluorescence intensity and frequent late amplification prevented reliable automatic Ct determination. Raw fluorescence data from cycles 1–40 were therefore exported and analyzed using a consistent procedure across all assays, instruments, and reporter dyes.

For each well, the mean fluorescence over cycles 5–15 was subtracted from all fluorescence measurements. For each dataset, the standard deviation (SD) of the baseline-subtracted fluorescence was calculated by pooling measurements from all 0-copy/µL control wells over cycles 5–15. The fluorescence threshold was set at 10 times this SD. Ct was determined from the first threshold crossing after cycle 15, using linear interpolation between adjacent cycles. No curve fitting was applied. Wells with Ct ≤ 38.0 were classified as detected; wells with Ct > 38.0 or no threshold crossing within 40 cycles were classified as not detected. Mic data were analyzed using the same procedure, with thresholds calculated from the corresponding Mic 0-copy/µL controls.

Standard curves were constructed by ordinary least-squares regression of Ct against the base-10 logarithm of input target concentration (copies/µL), using individual-well Ct values at all tested concentrations at or above the limit of detection (LOD). The LOD was operationally defined as the lowest tested concentration at which all replicate reactions were detected, provided that all higher tested concentrations were also fully detected. Amplification efficiency was calculated as E = (10^(−1/a) − 1) × 100%, where a is the regression slope. Standard-curve linearity was reported as the coefficient of determination (R²). Detection rates were calculated as the number of detected wells divided by the total number of replicate wells at each concentration. Replicate numbers are provided in the corresponding figure legends. For comparison between the Mic and QuantStudio platforms, Ct values at matched BDBV RNA concentrations were compared by linear regression. The regression slope and R² were reported to characterize the relationship between the two platforms.

## ACKNOWLEDGMENTS

OpenAI’s ChatGPT, Google’s Gemini, and Anthropic’s Claude were used to assist with language editing, sentence structure, readability, reference formatting, and the preparation of schematic illustrations. These AI tools were not used for experimental data generation or analysis. All AI-assisted content was reviewed and verified by the authors, who take full responsibility for the final manuscript.

## AUTHOR CONTRIBUTIONS

**J.L.:** Conceptualization (lead); Methodology (lead); Investigation (lead); Formal analysis (lead); Visualization (lead); Writing – original draft (lead).

**E.S., S.N., and R.T.:** Resources (supporting); Investigation (supporting); Methodology (supporting).

**S.B., K.M., T.A., B.H., D.N., N.E., R.M., H.K., and C.W.:** Investigation (supporting).

**A.O.:** Supervision (supporting); Writing – review & editing (supporting).

**P.C.S.:** Supervision (lead); Resources (lead); Funding acquisition (lead); Writing – review & editing (supporting).

All authors reviewed, edited, and approved the final manuscript.

## DATA AND CODE AVAILABILITY

All data supporting the findings of this study are publicly available in Zenodo at https://doi.org/10.5281/zenodo.23921512.

