## Supplementary Information for "Direct Whole-Blood PCR for Molecular Diagnostics"

**Table S1.** Scoring criteria used to evaluate diagnostic technologies according to the REASSURED framework. The 3-point scoring scheme was adapted and slightly modified from Otoo et al.[52]

| Criteria | High (3) | Medium (2) | Low (1) |
|---|---|---|---|
| **R – Real-time connectivity** | Test results are automatically interpreted and electronically transmitted to a connected or centralized system, enabling real-time data sharing, clinical decision-making, or surveillance. | Test results are automatically interpreted and reported by the device, but the system lacks integrated electronic connectivity for real-time remote data transmission. | Test results require manual interpretation by the user (e.g., visual interpretation of colorimetric or lateral-flow readouts) and are not automatically reported or transmitted. |
| **E – Ease of specimen collection** | Finger prick, sputum | | Venous blood, serum |
| **A – Affordable** | x < $5 | $5 < x < $20 | x > $20 |
| **S – Sensitive** | x > 95% | 90% < x < 95% | x < 90% |
| **S – Specific** | x > 95% | 90% < x < 95% | x < 90% |
| **U – User-friendly** | x < 2 minutes, CLIA waived | 2 < x < 5 minutes, moderate complexity | x > 5 minutes, high complexity |
| **R – Rapid and robust** | x < 1 hr | 1 < x < 3 hrs | x > 3 hrs |
| **E – Equipment-free/simple** | Portable, handheld, disposable device or cartridge, battery or solar powered. | Portable, disposable device or cartridge, that can potentially be powered by a power pack. | Portable device or cartridge, potentially powered by power pack. |
| **D – Deliverable** | Reagents can be stored at room temp | Reagents are stable at room temp but for a few hours | Reagents require refrigeration |

**Table S2.** Estimated per-reaction costs of reagents and consumables.

| Component | Cat No | Cost ($) | Number of reactions | Cost per run ($) |
|---|---|---|---|---|
| Q5 master mix | M0500L | 542 | 500 | 1.084 |
| Primer | | 10 | 2000 | 0.005 |
| Reporter | S11363 | 390 | 1250 | 0.312 |
| Reverse Transcriptase | E3005E | 2,549 | 2500 | 1.020 |
| Mic Tube | | 104 | 960 | 0.108 |
| Sum | | | | 2.529 |

**Table S3.** Comparison of direct whole-blood molecular diagnostic technologies.

| No. | Study / technology | Actual sample entering amplification | Treatment / processing | Strict direct WB? | WB input | Cost reported | Whole-blood analytical sensitivity / LOD | Clinical sensitivity | Clinical specificity | Hands-on / workflow | Time-to-result |
|---|---|---|---|---|---|---|---|---|---|---|---|
| 1 | **This Work** | Unrefined whole blood | DNA: none. RNA: RNase inhibitor added to WB at 1:39 v/v → directly into RT-qPCR | Yes | 2 µL / 20 µL = 10% | ~$2.53/reaction; portable device ~$17,000 | HBV: 10 copies/µL or 20 copies/reaction; HCV/BDBV/EBOV/SUDV: 100 copies/µL or 200 copies/reaction | NR – no clinical cohort | NR – no clinical cohort | <2 min sample-to-instrument | ~1 h |
| 2 | **Lim et al. 2026 – biphasic dried blood RT-LAMP**[20] | Dried whole-blood matrix | RNase inhibitor → 37°C 30 min drying → 95°C 10 min thermal lysis → 55°C 20 min RT → 65°C 1 h RT-LAMP | No | 4 µL | ~$3.59/reaction; portable reader ~$60 | Zika: 10 copies/µL; HCV: 1 IU/µL | NR – analytical study | NR – no clinical cohort | Multiple timed steps and reagent additions | ~2 h |
| 3 | **Semper et al. – GeneXpert Ebola**[53] | Starts with venous EDTA whole blood, but cartridge receives chemically treated sample | 100 µL WB + 2.5 mL sample reagent → 20 min inactivation → cartridge → automated sample preparation/extraction + RT-PCR | Yes | 100 µL | ~$20/test under subsidized/access pricing; | N/A | 100% (22/22; 95% CI 84.6–100%) | 95.8% primary analysis | Low operator burden after cartridge loading | ~2.5 h sample receipt → result |
| 4 | **Mehta et al. – direct DENV RT-PCR**[54] | Whole blood directly into RT-PCR for analytical testing; plasma for clinical cohort | No extraction or separate lysis. Direct Blood RT-PCR buffer + RNase inhibitor + high SYBR concentration | Yes, but based on melting curve analysis | 2.5 µL | Per-test cost NR; portable Open qPCR ~$4,299 | $1{\times}10^4$ GCE/mL = 10 GCE/µL in whole blood | Portable system: 78.3%; benchtop: 76.7% | Portable: 90.9%; benchtop: 93.9% | Relatively low; direct reaction setup followed by RT-PCR and melt analysis | ~1.5–2 h |
| 5 | **Shah et al. – QuRapID Ebola**[55] | Fresh whole blood directly into reaction for blood-compatibility workflow | Blood enters reaction directly, followed by 8–10 automated freeze–thaw cycles (~2 min) for virion disruption before RT-qPCR | No | 5 µL | $12.50/test disposables | Live EBOV: minimum detected $1.4{\times}10^3$ PFU/mL; LLOQ $1.4{\times}10^4$ PFU/mL | NR – no patient clinical cohort | NR – analytical specificity only | Freeze–thaw automated; otherwise minimized manual steps | ~70 min |
| 6 | **Cai et al. – direct nested PCR/RT-PCR**[56] | Anticoagulated whole blood directly into first amplification | No sample pretreatment, but first PCR/RT-PCR → tube opening/transfer → nested real-time PCR | Yes, but not single-step | 4–20 µL | NR | DNA: 5.8 copies/µL at 10–30% WB; RNA: $6.8{\times}10^3$ copies/µL; intact bacteria: 6.6 CFU/µL | NR – no clinical cohort | NR – analytical specificity only | High – two amplification rounds and open-tube transfer | <4 h |

| | | | | | | | | | | | |
|---|---|---|---|---|---|---|---|---|---|---|---|
| **7** | **Li et al. – ZIKV dirRT-qPCR**[57] | EDTA-K2 whole blood directly into RT-qPCR | No separate pretreatment; inhibitor-tolerant polymerase + PEC-2 / polymerase enhancers incorporated into amplification mixture | Yes | 5 µL | NR | Whole blood: $9.5\times10^2$ copies/reaction | 100% (9/9 ZIKV-positive clinical specimens) | 100% (74/74 negative specimens) | Relatively low; direct sample addition, no extraction | <2 h |

**Table S4.** Oligonucleotides used in this study

| Assay | Role | Sequence (5'→3') | Used in |
|---|---|---|---|
| HBV | Forward primer | GATGTGTCTGCGGCGTTTTA | Fig 2, 5, 6b–i |
| HBV | Reverse primer | GCAACATACCTTGATAGTCCAGAAGAA | Fig 2, 5, 6b–i |
| HCV | Forward primer | TGCACGGTCTACGAGAC | Fig 3, S1 |
| HCV | Reverse primer | GCCTTGTGGTACTGCCTGAT | Fig 3, S1 |
| BDBV | Forward primer | TAACTCATCCACAGACCCCA | Fig 4a–b, 6j–k |
| BDBV | Reverse primer | GATGGCGCCGAAACTATAGA | Fig 4a–b, 6j–k |
| EBOV | Forward primer | ACAAATTGGATGCTCTACATGT | Fig 4c–d |
| EBOV | Reverse primer | TCCACCAGAAAACCCATGTT | Fig 4c–d |
| SUDV | Forward primer | ACCGACATTTAGACACTCAAGT | Fig 4e–f |
| SUDV | Reverse primer | ATAATCTTGCAGCGTCCGAT | Fig 4e–f |
| HBV | Quantitative synthetic DNA standard (not an ordered oligo, ATCC VR-3232SD) | | Fig 2, 5, 6b–i |
| HCV | Quantitative synthetic RNA standard (not an ordered oligo, ATCC VR-3233SD) | | Fig 3, S1 |

| | | | |
|---|---|---|---|
| BDBV | Synthetic DNA template for in vitro transcription (T7 promoter + target region) | gaaatTAATACGACTCACTATAgggTACCTAACCTACACGTCTACGCTTTCCTTGGATCTCACAAGGTACCGAGAGAATGAGTTAATTTATGATAACAATCCGTTAAAAGGTGGACTTAATTGCAACCTATCCTTTGATAATCCACTTTTCAAGGGCCAAAGGCTCAATATCATAGAGGAGGATTTGATTAGATTTCCTCATCTATCTGGGTGGGAACTTGCGAAACCATCATTCAGTCCATTATCTCAGACAGCAATAACTCATCCACAGACCCATTAGCAGTGGAGAAACACGATCATTCACAACTCACTTTCTCACATATCCTAAGGTTGGGCTCCTCTATAGTTTCGGCGCCATCGTGAGTTATTACTTAGGGAATACCATTATTAGGACCAAAAAGCTAGACCTCAGTCATTTTATGTATTACTTAACAACTCAAATCCATAATTTGCCACATCGCTCGTTGAGGATACTTAAGCCCACCTTTAAACATGTTAGTGTGATATCAAGACTAATGAGTAT | Template for RNA standards (Fig 4, 6) |
| EBOV | Synthetic DNA template for in vitro transcription (T7 promoter + target region) | gaaatTAATACGACTCACTATAgggCAGCTGATTGACCAGTCTTTGATTGAGCCCTTAGCAGGAGCCCTTGGTCTGATCTCTGATTGGCTGCTAACAACCAACACTAACCATTTCAACATGCGAACACAACGTGTCAAGGAACAATTGAGCCTAAAAATGCTGTCGTTGATTCGATCCAATATTCTCAAGTTTATTAACAAATTGGATGCTCTACATGTCGTGAACTACAACGGATTGTTGAGCAGTATTGAAATTGGAACTCAAAATCATACAATCATCATAACTCGAACTAACATGGGTTTTCTGGTGGAGCTCCAAGAACCCGACAAATCGGCAATGAACCGCAAGACACCTGGGCCGGCGAAATTTTCCCTCCTTCATGAGTCTACACTGAAAGCATTTACACAAGGATCCTCGACACGAATGCAAAGTTTGATTCTTGAATTTAATAGCTCTCTTGCTATCTAACTAAGATGGAATACTTCATATTGAGCTAACTCATATATGCTGACTCAATAGTTATC | Template for RNA standards (Fig 4) |
| SUDV | Synthetic DNA template for in vitro transcription (T7 promoter + target region) | gaaatTAATACGACTCACTATAgggTTAGCAGCGGTGAAACAAGATCCTTCACAACCCACTTCTTAACGTATCCCAAAATAGGGCTCCTATACAGTTTTGGAGCCCTCATAAGTTTTATTTGGGTAATACTATTCTGTGCACGAAAAGATCGGACTCACAGAATTTCTATACTATCTCCAGAATCAGATCCACAACTTATCACACAGATCCCTTCGAATCTTCAAACCGACATTTAGACACTCAAGTGTCATGTCCAGGTTGATGGATATAGACCCCAACTTCTCAATATATTGGTGGGACTGCAGGTGACCGTGGATTATCGGACGCTGCAAGATTATTTCTCCGAATTGCAATTTCAACTTTCTTGAGCTTTGTTGAGGAGTGGGTTATCTTTAGGAAGGCAAACATCCCACTATGGGTTATCTATCCTCTCGAAGGCCAACGCCCTGATCCCTCTGGCGAATTTTTGAACCGAGTAAAATCTCTAATTATTGGGACTGAAGATGATAAAATAAAGGT | Template for RNA standards (Fig 4) |

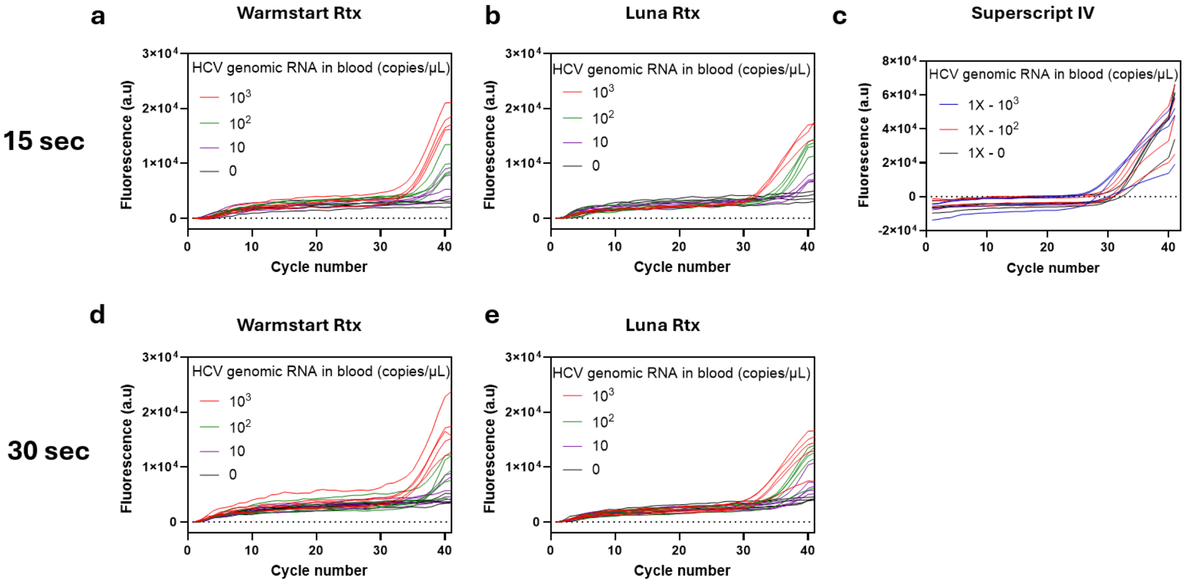

**Figure S1. Optimization of reverse transcriptase and extension time for direct whole-blood RT-qPCR.** Amplification profiles of HCV synthetic RNA directly spiked into whole blood using different reverse transcriptases and extension times. Reverse transcription was performed at 55 °C for 10 min, followed by qPCR with extension times of either 15 or 30 s. (a–c) Amplification profiles obtained with a 15-s extension using (a) WarmStart® RTx Reverse Transcriptase, (b) Luna® WarmStart® RT Enzyme Mix, and (c) SuperScript™ IV Reverse Transcriptase. (d, e) Amplification profiles obtained with a 30-s extension using (d) WarmStart® RTx Reverse Transcriptase and (e) Luna® WarmStart® RT Enzyme Mix. HCV RNA concentrations of $10^3$, $10^2$, 10, and 0 copies/μL were evaluated. Luna® WarmStart® RT Enzyme Mix with a 15-s extension (b) demonstrated the most distinct concentration-dependent amplification profiles, whereas SuperScript™ IV (c) exhibited nonspecific amplification in the whole-blood matrix. Each curve represents an individual reaction (n = 4–5 per condition).

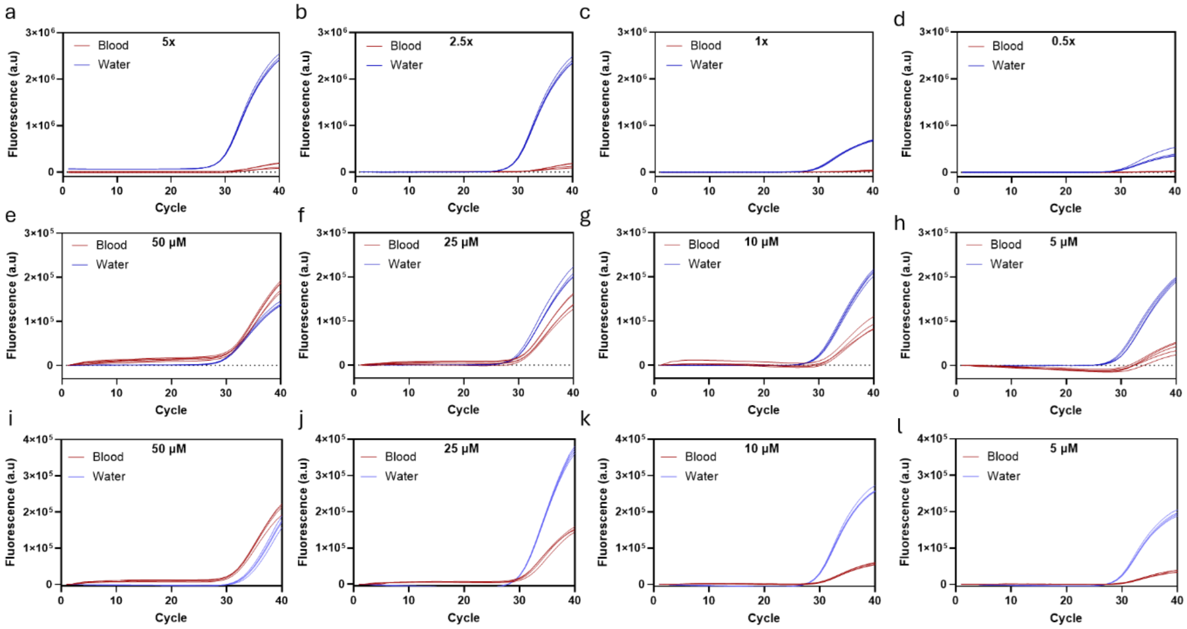

**Figure S2.** Raw fluorescence data comparing the HBV direct qPCR assay in whole blood and water at different dye concentrations: (a–d) EvaGreen, (e–h) SYTO 82, and (i–l) SYTO 61.

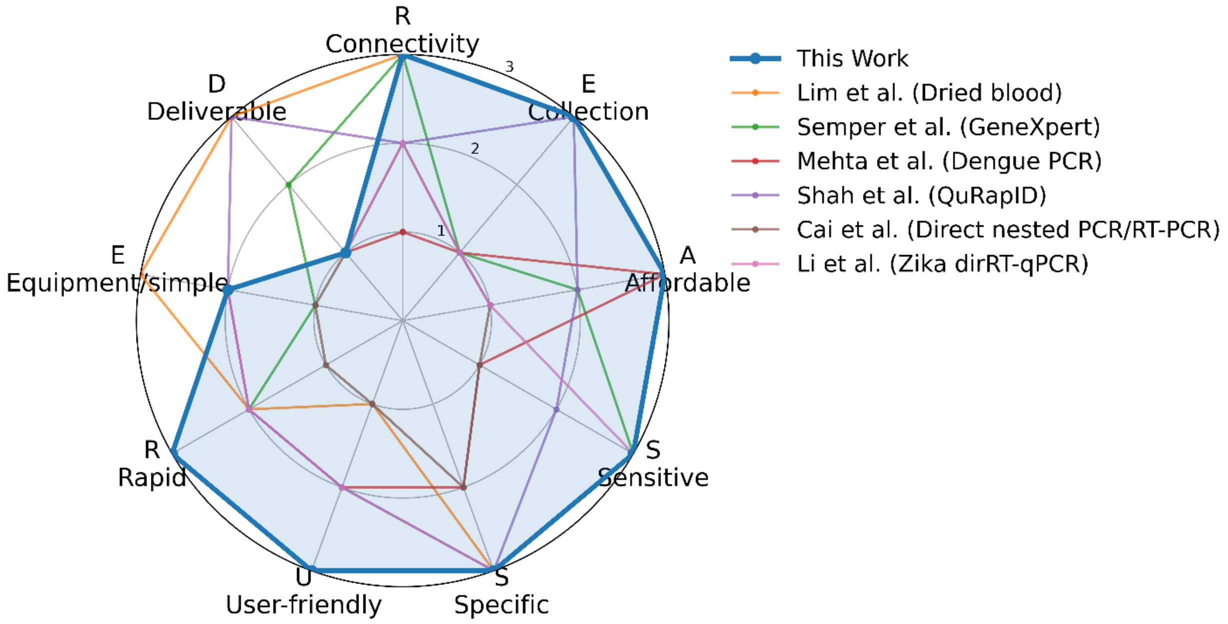

| Diagnostic (reference) | R | E | A | S | S | U | R | E | D | Total (/27) |
|---|---|---|---|---|---|---|---|---|---|---|
| **This work**<br>**Direct whole-blood RT-qPCR** | 3 | 3 | 3 | 3 | 3 | 3 | 3 | 2 | 1 | **24 (89%)** |
| Lim et al. (Dried blood)<br>Direct dried-blood RT-LAMP | 3 | 3 | 3 | 3 | 3 | 1 | 2 | 3 | 3 | **24 (89%)** |
| Semper et al.<br>GeneXpert Ebola | 3 | 1 | 2 | 3 | 3 | 2 | 2 | 1 | 2 | **19 (70%)** |
| Mehta et al.<br>Direct Dengue PCR | 1 | 1 | 3 | 1 | 2 | 2 | 2 | 2 | 1 | **15 (56%)** |
| Shah et al.<br>QuRapID Ebola | 2 | 3 | 2 | 2 | 3 | 2 | 2 | 2 | 3 | **21 (78%)** |
| Cai et al.<br>Direct nested PCR/RT-PCR | 2 | 1 | 1 | 1 | 2 | 1 | 1 | 1 | 1 | **11 (41%)** |
| Li et al.<br>Direct Zika RT-qPCR | 2 | 1 | 1 | 3 | 3 | 2 | 2 | 2 | 1 | **17 (63%)** |

**Figure S3.** REASSURED assessment of representative direct whole-blood molecular diagnostic technologies.

### SUPPLEMENTARY NOTES

#### Supplementary Note 1. Practical benefits and remaining gaps toward a REASSRUED diagnostic.

**Benefits of direct whole-blood amplification.** Direct testing of a small blood volume could provide four simplifications: no delivery, no large-volume sampling, no sample preparation, and no operational difficulty.[58,59] Testing at the site of collection could eliminate specimen delivery and thereby reduce transport-associated cold-chain requirements,[60,61] sample degradation,[62,63] and anticoagulant considerations associated with storage and transport.[64,65] Small-volume detection could avoid large-volume venous blood collection and the excess specimen volume that downstream extraction requires; we did not evaluate finger-prick sampling. Eliminating sample preparation could reduce reagents, consumables, instrumentation, sample transfers, and operator steps,[60,66] and sufficiently simple, closed operation could reduce training requirements.[66]

The aim is to combine the operational simplicity of the rapid diagnostic tests and lateral-flow assays widely used during the COVID-19 pandemic with an analytical sensitivity that approaches standard molecular methods. If achieved, this combination could shorten diagnostic delay and reduce the need for patients to revisit healthcare facilities, supporting timely clinical intervention for individual patients and faster case identification and containment at the public-health level.

**Remaining REASSURED gaps (Fig. S3).** The gaps below are engineering and formulation tasks. They are separate from the scientific questions addressed in the Discussion, namely performance with intact pathogens and in patient specimens, which must be answered first.

**Connectivity and power.** Real-time connectivity depends mainly on access to wireless or internet infrastructure, which continues to expand through cellular and satellite-based communication.[67] The Mic instrument operates on AC power, and the manufacturer reports that it can run in the field from a battery or solar source through a pure sine wave inverter (360 W minimum). The inverter adds equipment, and an instrument with integrated battery operation would simplify deployment. Additionally, portable fluorescence detection has been demonstrated with compact, iPod-sized instruments, including the VPodDuo system.[20,68] Equipment-free or minimally instrumented operation may therefore benefit from advances in low-power electronics, optical detection, and device miniaturization.

**Deliverability.** Deliverability depends on reagent formulation. Lyophilized reagents are widely implemented for PCR-,[69] LAMP-,[16,20] and RPA-based[19] assays. The NEB Q5® Blood Direct Master Mix used here contains proprietary components that enable amplification in the presence of whole-blood inhibitors, and it is not currently available in a lyophilized format. For the RNase inhibitor required for RNA detection, glycerol-free formulations should facilitate lyophilization.[70]

#### Supplementary Note 2. Cartridge-based diagnostics and the cost basis of the comparison

Cartridge-based systems such as the GeneXpert Ebola assay achieve an analytical sensitivity of approximately 230 copies/mL by miniaturizing centralized laboratory workflows into complex microfluidic consumables.[50] Adapting a laboratory workflow, rather than designing an assay for the point of care, inherently increases cost.[20] Public-sector subsidies may reduce the cartridge price to approximately $20 per test in some regions,[71] whereas the unsubsidized average price charged by private laboratories or hospitals often ranges from $68 to more than $150.[72]

We estimated the reagent and consumable cost of our workflow at approximately $2.53 per reaction (Table S2), using commercially available components without bulk purchasing or manufacturing-scale optimization. Larger-scale procurement may reduce this cost further. This figure covers reagents and consumables only. It excludes the instrument (Table S3 lists approximately $17,000 for the portable device) and labor, and it is a cost per reaction rather than a price per test, so it is not directly comparable with cartridge prices.

High-cost cartridge-based models may be difficult to sustain at scale in resource-limited settings. They can perpetuate a reactive cycle in which each new outbreak requires renewed governmental or international intervention to finance and deploy diagnostic capacity, rather than testing infrastructure that is accessible locally before an outbreak emerges.[73]

Among direct whole-blood assays, the Zika RT-qPCR assay was evaluated in clinical specimens, with 100% sensitivity in 9/9 ZIKV-positive specimens and 100% specificity in 74/74 negative specimens (Table S3).[57] Its detection limit was approximately one order of magnitude higher than ours. We did not test patient specimens, so we do not report clinical sensitivity or specificity for our assays (Table S3).

#### Supplementary Note 3. Sample-volume basis for comparing direct and extraction-based assays.

Detection limits expressed in copies/µL of sample can be converted to copies/mL; however, unit conversion alone does not allow direct comparison between direct and extraction-based assays because the two workflows analyze different effective sample volumes and involve different sample processing steps. The difference arises from the scaling limits of nucleic acid extraction. Solid-phase binding and elution on silica membranes or magnetic beads cannot be scaled efficiently to microliter inputs.[74] Automated platforms therefore process larger sample volumes (e.g., 100–200 µL) with multi-milliliter volumes of lysis and inactivation buffer (e.g., 2.5 mL of sample reagent for GeneXpert)[50] and elute into a smaller final volume (typically ~50 µL). Because no more than 5 µL of the purified extract enters the RT-PCR reaction, most of the processed material is never analyzed.

This upfront volumetric concentration favors extraction-based assays when sensitivity is expressed per milliliter. For example, a direct PCR assay with an LOD of 100 copies/µL and a 2-µL whole-blood input starts amplification with 200 target copies per reaction. An extraction-based assay that processes 100 µL of sample and reports an LOD of 10 copies/µL of sample starts with 1,000 target copies. Extrapolating the direct-PCR LOD from copies/µL to copies/mL therefore ignores the absence of upfront volumetric enrichment and does not give an equivalent metric. We compare detection limits in the units each study reported (Table S3).

69. LyoPrime Luna® Probe One-Step RT-qPCR Mix with UDG | NEB. https://www.neb.com/en-us/products/l4001-lyoprime-luna-probe-one-step-rt-qpcr-mix-with-udg?srsltid=AU7gw4XDTqJVh1J466nNCWKMfRgkHe8DKsHzKSWKheDAoKzu4w8A70S1.

70. RNase Inhibitor, Murine (Glycerol-free) | NEB. https://www.neb.com/en-us/products/m0628-rnase-inhibitor-murine-glycerol-free?srsltid=AU7gw4X9E9qYMkCZLYLethTHmeeOyFBgVRGUQdaa84B-iKM9CUIhLksl.

71. Raftery, P. *et al.* Establishing Ebola Virus Disease (EVD) diagnostics using GeneXpert technology at a mobile laboratory in Liberia: Impact on outbreak response, case management and laboratory systems strengthening. *PLoS Negl Trop Dis* **12**, e0006135 (2018).

72. Ponnudurai, N., Denkinger, C. M., Gemert, W. V. & Pai, M. New TB Tools Need to be Affordable in the Private Sector: The Case Study of Xpert MTB/RIF. *Journal of Epidemiology and Global Health* **8**, 103–105 (2018).

73. The Lancet Regional Health – Western Pacific. Breaking the cycle of panic and neglect: better preparedness for the next pandemic. *The Lancet Regional Health - Western Pacific* **58**, 101586 (2025).

74. Obino, D. *et al.* An Overview on Microfluidic Systems for Nucleic Acids Extraction from Human Raw Samples. *Sensors* **21**, 3058 (2021).
